# Continent-wide assessment of the strain-level diversity of *Bradyrhizobium*, a dominant soil bacterial genus

**DOI:** 10.1101/2025.05.29.656829

**Authors:** Clifton P. Bueno de Mesquita, Matthew R. Olm, Andrew Bissett, Noah Fierer

**Author notes:** Corresponding authors: CP Bueno de Mesquita,; Noah Fierer.

## Abstract

Global surveys of soil bacteria have identified several taxa that are nearly ubiquitous and often the most abundant members of soil bacterial communities. However, it remains unclear why these taxa are so dominant across a wide range of soil types and environmental conditions. Here we use genome-resolved metagenomics to test the hypothesis that strain-level differences exist in these taxa that are not adequately captured with standard marker gene sequencing, and that distinct strains harbor unique traits that reflect adaptations to different soil environments. We analyzed data from 331 natural soils spanning Australia to assess strain differentiation in *Bradyrhizobium*, a dominant soil bacterial genus of ecological importance. We developed a workflow for strain-level bacterial analyses of complex soil metagenomes, combining genomes from pre-existing databases with new genomes generated via targeted assembly from metagenomes to detect 181 *Bradyrhizobium* strains across the soil collection. In addition to a high degree of phylogenetic variation, we observed substantial variation in pangenome content and inferred traits, highlighting the breadth of diversity within this widespread genus. While members of the genus *Bradyrhizobium* were detected in > 80% of samples, most individual strains were restricted in their distributions. The overall strain-level community composition of *Bradyrhizobium* varied significantly across geographic space and environmental gradients, and was particularly associated with differences in temperature, soil pH, and soil nitrate and metal concentrations. Our work provides a general framework and methodology for studying the strain-level ecology of soil bacteria and highlights the ecological and pangenomic diversity within this dominant soil bacterial genus.

## Introduction

Soils harbor some of the most diverse bacterial communities on Earth [1]. As in nearly all biological communities, most soil bacterial taxa are rare and only a few are abundant in any particular sample [2–4]. Moreover, the bacterial taxa that are abundant in any individual soil often tend to be abundant and ubiquitous across many different soils that span a range in environmental conditions [2, 5–7]. This observation then raises the question of why certain bacterial taxa are so ubiquitous and abundant in soil. How are these ‘dominant’ taxa able to thrive in vastly different soil environments? The most parsimonious explanation is there are species- to subspecies-level differences within a given taxon across different environments, with high strain-level diversity corresponding to high trait diversity, such that members of a single taxon can persist across a range of environmental conditions [8]. In other words, a bacterial genus or species that appears to be dominant across many soils is actually composed of many distinct strains with distinct ecologies.

There is a growing body of research in both environmental and host-associated systems highlighting the value of strain-level analyses for exploring the structure of bacterial communities and identifying spatiotemporal patterns that may not be apparent at coarser levels of taxonomic resolution [9]. For example, Cho and Tiedje (2000) observed endemism of different *Pseudomonas* species in different locations, suggesting that the species within a ubiquitous and abundant genus were not globally mixed [10]. More recently, Coleman and Chisholm (2010) found that there was a large fraction of genes that were rare in populations of two abundant marine bacteria, reflecting continual gene transfer and loss [11]. Furthermore, a pangenomic study of *Prochloroccus,* one of the most ubiquitous marine bacterial genera, revealed that subtle differences in gene cluster content were associated with biogeographic patterns [12]. Even in the human microbiome, recent work has shown how strain-level analyses can be used to understand bacterial community assembly in infant guts [13, 14].

Shotgun metagenomic sequencing and subsequent genome-resolved analyses can help refine taxonomic identifications and functional gene analyses to the species and strain levels of resolution [15]. There are many tools for such strain-based analyses of bacteria using metagenomic data, including reference-based and *de novo* approaches, with such tools frequently used to investigate the human gut microbiome where the reference genome databases are reasonably comprehensive [16–18]. Performing strain-level analyses on environmental samples, particularly soils, is far more challenging due to the complexity of these communities and the paucity of reference genomes available for many members of these communities, even the more abundant taxa [19]. In recent years, reference genome databases have grown exponentially in size and the greater sequencing depth that can be achieved with newer sequencing technologies has enabled more effective analyses of complex environmental microbiomes. Still, given the reference genome database limitations, it is important to supplement those databases with genomes assembled directly from metagenomic samples of interest. While strain-level analyses of such *de novo* metagenome-assembled genomes (MAGs) have been the focus of previous work in soils and seawater [20, 21], here we develop a general methodological framework that combines available reference genome data with targeted assembly of additional genomes directly from soil metagenomes (Figure 1).

**Figure 1.**
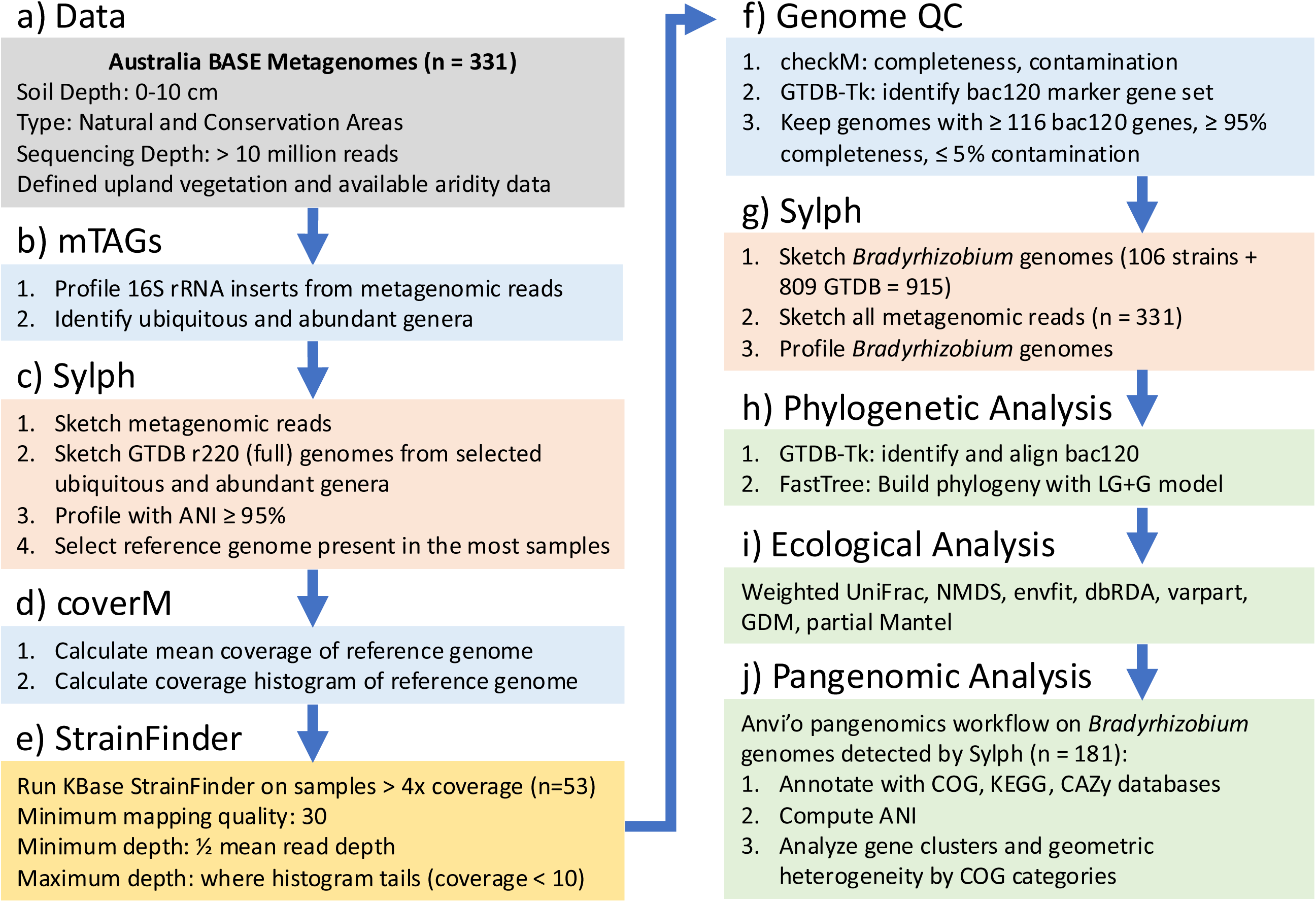
Methodological flow diagram including the starting data, identification of ubiquitous and abundant genera, selection of reference genomes, coverage calculation, targeted genome assembly with StrainFinder, genome QC, identification and relative abundance calculation (profile) with Sylph, and downstream phylogenetic, ecological, and pangenomic analyses.

We used this approach to investigate the strain-level diversity of the abundant and ubiquitous soil bacterial genus *Bradyrhizobium* using a standardized collection of soil metagenomes from across Australia. *Bradyrhizobium* has consistently been found to be one of the most ubiquitous and abundant members of soil bacterial communities in large-scale 16S rRNA gene sequencing efforts [2, 5–7, 22]. For example, in an analysis of 16S rRNA gene data from 275 Australian soils, four of the top five most abundant amplicon sequence variants (ASVs) were members of the *Bradyrhizobium* genus; these ASVs were also among the most prevalent ASVs, being detected in > 74% of samples [6]. *Bradyrhizobium* are most well known as taxa that symbiotically fix nitrogen (N) with various plant hosts in the legume family and thus play an important role in N cycling and ecosystem productivity [23, 24]. *Bradyrhizobium* spp. containing a genomic island of *nif* genes for nitrogen fixation were found to be widespread in a variety of environments [25]. Indeed, *Bradyrhizobium* spp. are the dominant diazotrophs in many soils, often comprising over 50% of the N-fixing soil bacterial community [26]. However, the genus *Bradyrhizobium* also contains free-living soil-inhabiting taxa and taxa capable of photosynthesis [27–29]. While long known as symbionts of both legumes and non-legumes [23, 24], more recent work has highlighted the importance of non-symbiotic *Bradyrhizobium* in soils, as non-symbiotic members of the genus were found to dominate North American forest soils [29]. Given the dominance of the *Bradyrhizobium* in many soil bacterial communities and given the genomic and ecological diversity represented within this genus, we expected that we could use our approach (Figure 1) to conduct genome-resolved analyses of this genus and provide insights into the biogeographical patterns exhibited by this taxon that could not be resolved using more standard marker gene sequencing approaches.

We conducted strain-level genomic analyses of species within the genus *Bradyrhizobium* to investigate the biogeography and pangenome of this genus directly from 331 shotgun metagenomes representing a broad range of soil types from across the continent of Australia. We used the resulting data from our approach to test three hypotheses. First, no single strain of this genus is dominant in soils across a wide range of sites and environmental conditions, but rather there is extensive strain-level diversity within the genus, with specific strains having more restricted distributions. Second, both geographic distance and environmental differences are associated with *Bradyrhizobium* strain-level community composition. Finally, the detected *Bradyrhizobium* strains collectively have a large pangenome that reflects extensive ecological diversification within the genus.

## Materials and Methods

### Data

Quality-filtered shotgun metagenomic data (150 bp reads) were downloaded from the Australian BASE program via a request in their online web portal [30]. We selected 331 soil samples that met the following criteria: 0-10 cm depth, from natural areas (designated “natural and conservation areas” in BASE), defined upland vegetation types, sequencing depth of at least 10 million reads, and available data on climate characteristics (see below) (Figure 1a). We downloaded all associated sample and environmental metadata from BASE, which included edaphic properties such as organic carbon content, nitrate concentration, pH, available phosphorus, and other biogeochemical variables [30] (Figure S1). To complement the soil data, we acquired climate data from publicly available sources. Aridity index was determined for each sample using the Global Aridity Database version 3 map at 30 arc-second resolution [31], while mean annual temperature and precipitation were determined using the WorldClim2 database [32]. The 331 selected samples were distributed across most of the Australian mainland and the island of Tasmania and spanned large gradients in both climate (including arid, semi-arid, dry sub-humid, and humid climate classifications, [31]) and soil properties (Figure 2).

**Figure 2.**
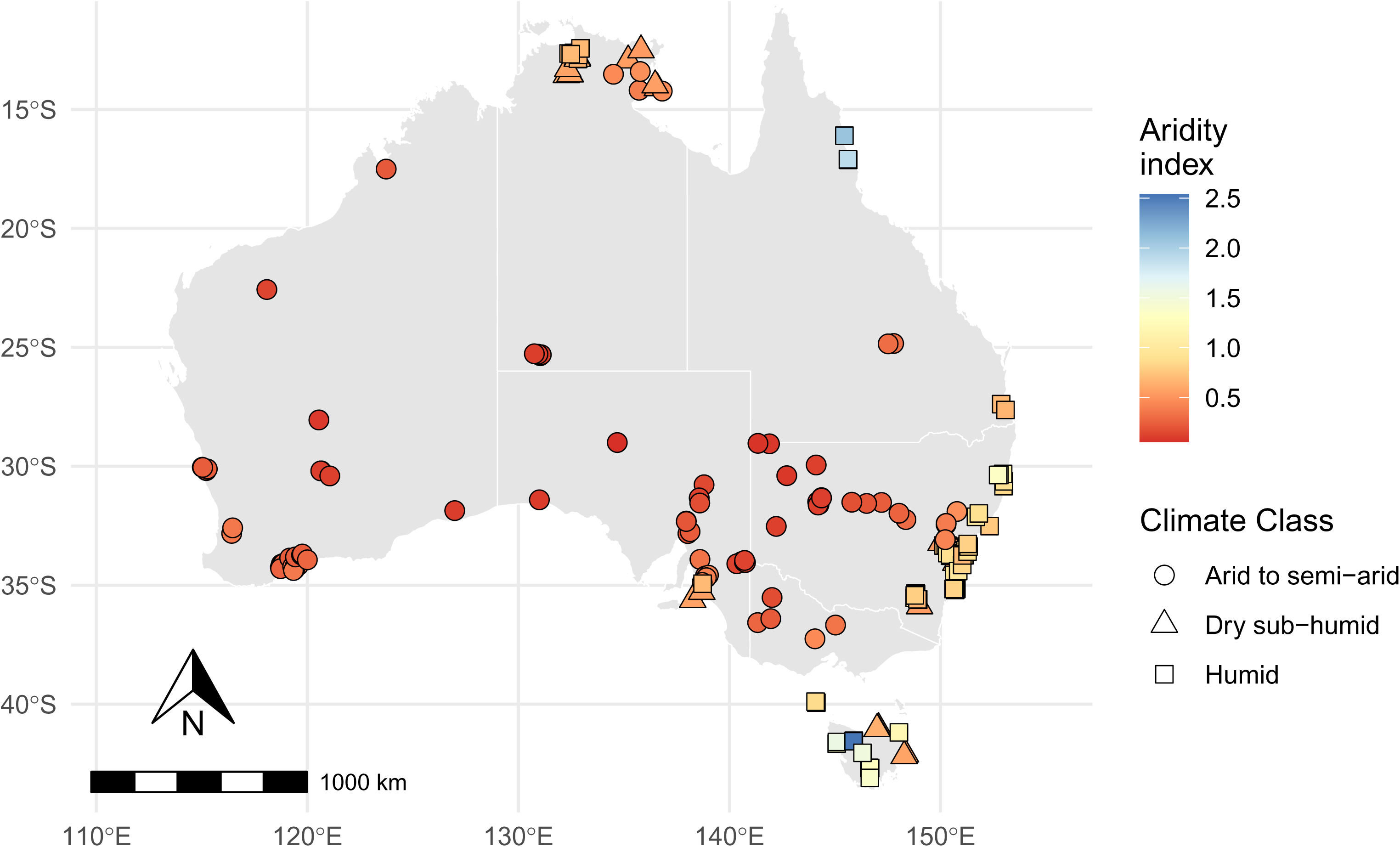
Map of the 331 BASE soil shotgun metagenomes used in this study with the sampling locations colored by their aridity index, where a higher aridity index indicates wetter conditions. These 331 samples include 103 “arid to semi-arid” samples, 110 “dry sub-humid” samples, and 118 “humid” samples.

### Genome Selection and Targeted Assembly

To confirm the abundance and ubiquity of *Bradyrhizobium*, we first selected a subset of 104 samples spanning most of the aridity gradient. We profiled the raw metagenomic reads taxonomically using mTAGs [33], which identifies 16S and 18S rRNA gene sequences in the metagenomes and assigns taxonomy by aligning them to the SILVA v138.1 database [34] with vsearch [35] (Figure 1b). We then downloaded all of the genomes of ubiquitous (75% of samples) and abundant (mean > 0.1% relative abundance across the whole dataset) genera from the full version of GTDB r220 [36] by first downloading the GTDB metadata to get the NCBI genome accessions for those genera, and then using NCBI datasets toolkit [37] to download the genomes (n = 21467) by accession. Next, to identify the best reference genomes for read mapping, we used Sylph [38] to build sketches of the 104 metagenomes and 21467 genomes, and determine which genomes were present using ‘profile’ mode and a 95% ANI cutoff (Figure 1c). To calculate read coverage and relative abundance, we then used coverM [39] to map reads to the most prevalent genomes (Figure 1d). A *Bradyrhizobium* genome (*Bradyrhizobium diazoefficiens_F*, GCA_016616885.1) was detected with mean coverage > 4x in the most samples (n = 53 of 104). High coverage is crucial for confident single nucleotide variant (SNV) calling and strain identification. We ran coverM with “method = mean” to calculate mean coverage and “method = coverage_histogram” to generate the histogram of coverage per base. Both methods were used to inform parameters in StrainFinder (see below) (Figure 1e).

The goal of using StrainFinder was to perform targeted assembly of *Bradyrhizobium* MAGs to capture diversity that is not captured within pre-existing reference genome databases. Strains were assembled with the KBase [40] implementation of StrainFinder [41], which maps reads to reference genomes, calls SNVs, and identifies strains based on SNV frequencies. We only profiled samples with mean coverage > 4x and at least 40% of the genome covered (n = 53). The KBase implementation then performs the additional steps of remapping reads to the consensus strains and assembling complete genomes for each strain. We used a default value of 30 for the read mapping quality parameter. We used the mean coverage output from coverM for each sample to define the minimum depth parameter for SNV calling in StrainFinder, which was set to half of the mean read depth. To avoid determining SNVs in regions that represent repeats, we set the maximum depth parameter to before the tail end of the coverage histogram for each sample. We defined the beginning of the “tail” of the coverage histogram as the coverage value at which the number of bases with that value was below 10. By default, four strains per sample were computed. We then used FastANI [42] to calculate the average nucleotide identity (ANI) among the strains and between the strains and the reference genome. Completeness and contamination of each strain were calculated with checkM [43].

To build a *Bradyrhizobium* genome database, we combined the 106 MAGs from StrainFinder (the top two strains per sample) with the 869 *Bradyrhizobium* genomes in GTDB r220, for a total of 975 *Bradyrhizobium* genomes. We used GTDB-tk [44] to identify a set of 120 single copy marker genes from those genomes (bac120). We filtered out any genome that was missing more than 4 of the bac120 genes, which yielded 915 genomes. The StrainFinder MAGs all had completeness > 95% and contamination < 1% and the GTDB genomes all had completeness > 95% and contamination < 5% (Figure 1f). These 915 genomes also included 19 commercial *Bradyrhizobium* inoculants used in Australian agriculture [45]. We used Sylph to sketch these 915 genomes and create a strain-by-sample relative abundance matrix across all 331 metagenomes using ‘profile’ mode and an ANI detection cutoff of 95% (Figure 1g).

### Phylogeny and Pangenomics

To build a phylogenetic tree of the starting 915 genomes and the Sylph-detected genomes (n = 181, see Results), we used GTDB-tk to align the previously identified bac120 genes. We used *Nitrobacter winogradskyi* Nb-255 (GCF_000012725.1) [46] as an outgroup to root the tree. Instead of randomly selecting 42 amino acids per gene (default setting), we initially included all amino acids per gene, but to minimize the effects of gaps, amino acids were retained only if they were present in 99% and 100% of the genomes for the 915 taxa tree and the 181 taxa tree, respectively. This yielded alignments of 2409 and 2026 amino acids, respectively. FastTree [47] was used to build the phylogeny using the LG+G substitution model (Figure 1h). The trees were plotted with the *ggtree* R package [48]. Pangenomic analysis of these 181 genomes was performed with anvi’o v8 [12, 49], which included functional annotation with the COG [50], KEGG [51], and CAZy [52] databases, ANI calculation with FastANI [42], computation of homologous gene clusters, and calculation of the geometric homogeneity index of each gene cluster [49] (Figure 1j). Geometric homogeneity compares the positions of gaps in the aligned residues without considering amino acid properties. To complement the ANI analysis, we also calculated the percent identity of full length 16S rRNA genes in the genomes, which were extracted with barrnap [53]. Estimated genome size was calculated by multiplying the assembly size by 100 and dividing by the checkM completeness percentage. Strains were classified as ‘symbiotic N-fixers’ if they contained at least five of six key *nif* genes (*nifBDEHKN*) and at least four of five key nodulation genes (*nodABCIJ*) [25], ‘free-living N-fixers’ if they only contained the *nif* genes and no *nod* genes, and ‘photosynthetic’ if they contained the photosynthetic reaction center HML subunits. We also searched for a suite of genes involved in aerobic respiration, carbon fixation, denitrification, single carbon compound (carbon monoxide, methane, methanol) metabolism, and sulfur metabolism [54].

### Ecological Analyses

The output of the Sylph ‘profile’ analysis was the relative abundance of each *Bradyrhizobium* strain (relative only to the other *Bradyrhizobium* strains in the sample) in each of the 331 soil metagenomic samples. We used the *pheatmap* R package to plot this relative abundance matrix along with relevant metadata [55]. We calculated a weighted UniFrac distance matrix in the *phyloseq* R package [56, 57]. Weighted UniFrac distance was significantly positively correlated with Bray-Curtis dissimilarity (Mantel r = 0.23, p < 0.001), but we opted to use weighted UniFrac because the Bray-Curtis dissimilarity matrix had a skewed distribution with many “1” values due to lack of any shared taxa among many samples, whereas weighted UniFrac accounts for phylogenetic similarity in the distribution of the *Bradyrhizobium* strains across samples and offers a metric for comparing two samples with no overlapping strains. Weighted UniFrac distance was also significantly positively correlated with unweighted UniFrac distance (Mantel r = 0.6, p < 0.001). To include more predictor variables for certain analyses, we subset the data to 219 samples that had a suite of 13 soil chemistry variables available. We performed non-metric multidimensional scaling (NMDS) combined with ‘envfit’ in the *vegan* R package [58] to visualize compositional differences and associations with environmental variables. We used distance-based redundancy analysis (dbRDA) and forward stepwise model selection in *vegan* to identify the top drivers of compositional differences. We partitioned variation attributed to geographic and dbRDA-selected environmental variables using ‘varpart’ in *vegan*. We used partial Mantel tests in *vegan* to assess correlations between weighted UniFrac distances and geographic distance and environmental distance (Euclidean distance of dbRDA-selected variables). We also used generalized dissimilarity modeling in the *gdm* R package [59] to further assess the effects of geographic distance and environmental dissimilarity and assess their relative importance (Figure 1i). All plots besides the heatmap were made with the *ggplot2* R package [60]. All analyses were performed with R version 4.2.3 [61]. Downstream data and analyses are publicly available on Zenodo (DOI: 10.5281/zenodo.15540063).

## Results

We first aimed to establish a comprehensive *Bradyrhizobium* genome database. Although *Bradyrhizobium* genomes are reasonably well-represented in reference databases (869 genomes in GTDB r220), we were concerned that the pre-existing genomes may not necessarily represent the full extent of *Bradyrhizobium* diversity in soil, especially since most (∼90%) of the pre-existing *Bradyrhizobium* genomes in GTDB are derived from cultured isolates and many members of this genus have not yet been cultivated [62]. Thus, to expand the diversity of *Bradyrhizobium* genomes used for our strain-level analyses, we implemented a StrainFinder-based pipeline for the targeted assembly of *Bradyrhizobium* genomes from a subset of the Australian soil metagenomes. StrainFinder generated 106 high-quality *Bradyrhizobium* MAGs (two from each of 53 samples). Estimated completeness of those strains ranged from 98.8% to 100% while contamination ranged from 0% to 0.34% (Figure S2). Average nucleotide identities (ANI) across the 106 strains ranged from 92 to 97%, and the ANI between the strains and the reference genome used in StrainFinder ranged from 93% to 97%. The percent identity of the full length 16S rRNA gene among the 106 StrainFinder MAGs ranged from 87.73% to 99.73%, and between the strains and the reference genome from 90.54% to 97.12%. All 106 StrainFinder MAGs were classified as genus *Bradyrhizobium* by GTDB-Tk, and the concatenated bac120 phylogenetic tree placed them in the *Bradyrhizobium* genus (Figure S3).

We next combined the 106 MAGs generated as described above with 809 high-quality pre-existing *Bradyrhizobium* genomes from GTDB and used Sylph [38] to determine which *Bradyrhizobium* genomes were detected in each of the 331 soil metagenomes (Figure 2). A total of 181 unique strains were detected (92 StrainFinder MAGs and 89 GTDB genomes) across 268 soils (81% of the soil metagenomes) (Figure 3, Figure S4). Of the 89 detected GTDB genomes, only 26 were from isolates, while a majority (63 genomes) were MAGs. The phylogeny of these 181 strains based on bac120 showed two clusters of GTDB genomes and a cluster of StrainFinder MAGs (Figure 3). These 181 strains were distributed across the broader *Bradyrhizobium* phylogeny (Figure S3). The strains detected included 6 of 19 commercial *Bradyrhizobium* strains [45]. All 53 of the most abundant StrainFinder MAGs were detected, while 39 of the 53 second most abundant StrainFinder MAGs were detected. The maximum number of strains detected per sample was 7, while the maximum number of samples a particular strain was detected in was 41. The estimated genome sizes for the 181 strains detected by Sylph ranged from ∼7 to 12 MB. Prevalence (number of samples in which a given strain was detected) was not significantly associated with genome size (linear regression, R^2^ = 0.02, p = 0.06, Figure S5) nor did genome size significantly differ between N-fixing and non-N-fixing *Bradyrhizobium* genomes (t-test, t = −0.66, p = 0.52). Patterns in the occurrence of strains in soils based on either aridity or mean annual temperature did not closely follow the phylogeny, but there was a high degree of variation in the range of temperatures the strains were detected in, with the widest range being 20.1°C (Figure S6, Figure S7). However, both aridity range and temperature range (for strains found in at least two soils) was more related to prevalence (linear regression, R^2^ = 0.34, p < 0.001 ; R^2^ = 0.30, p < 0.001, respectively) than to genome size (linear regression, R^2^ = 0.01, p = 0.52 ; R^2^ = 0.04, p = 0.08, respectively).

**Figure 3.**
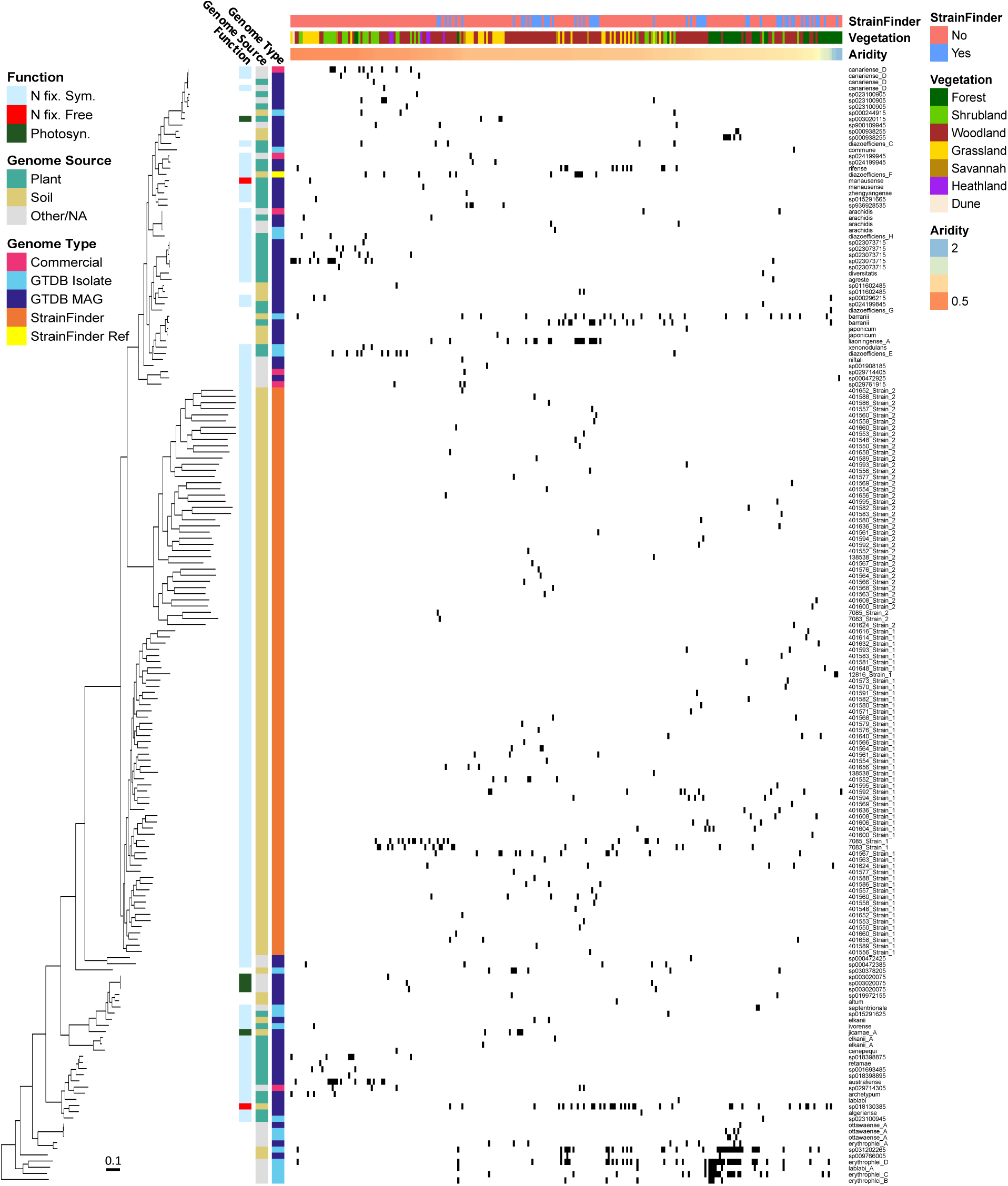
Presence (black) of 181 *Bradyrhizobium* strains detected by Sylph run on 331 samples and 915 *Bradyrhizobium* genomes (809 GTDB genomes + 106 StrainFinder MAGs). Shown are 181 strains (89 GTDB + 92 StrainFinder) detected in at least one of the 331 samples (in only 268 of the soil metagenomes did we detect at least one *Bradyrhizobium* strain). Columns (samples) are sorted by increasing aridity index from left to right while rows (genomes) are sorted according to the phylogenetic tree shown on the left, based on a concatenated alignment of 2026 amino acids from the bac120 marker gene set. Shown above the heatmap (legend on right) are sample metadata (whether or not StrainFinder was run on the sample, vegetation type, and aridity index). Shown to the left of the heatmap (legend on left) are genome metadata (function, genome source, and genome type). Genome source “Other/NA” means that the source was not stated in the metadata. For relative abundances of the strains in each soil metagenome, see Figure S4.

Among those 181 strains detected across the Australian soils, ANI ranged from 79.6% to 99.9% and full-length 16S rRNA gene percent identity (calculated for 179 strains that contained a full length 16S rRNA gene) ranging from 88% to 100%. Full length 16S rRNA gene percent identity among the most abundant StrainFinder MAGs and the GTDB genomes ranged from 92.76% to 100%, and among just the 89 detected GTDB genomes from 95.77 to 100%. These 181 strains would correspond to 82 operational taxonomic units (OTUs) at 99% similarity for the full-length 16S rRNA gene [63], 38 OTUs at 97% similarity for the V4 region of the 16 rRNA gene [64, 65], or 104 “species” at the 95% ANI cut-off [42, 66] (Figure S8). ANI calculations were not affected by plasmid sequences; of the 181 genome assemblies, only two had assembled plasmids and these plasmids were < 1 MB in size.

*Bradyrhizobium* community composition, as determined by the relative abundances of the 181 strains in the 268 soil metagenomes, was significantly associated with several soil and climate variables (Figure S9, envfit, p = 0.001). Distance-based redundancy analysis (dbRDA) identified temperature, pH, manganese, aluminum, nitrate, and zinc as the most significant predictors of *Bradyrhizobium* community composition (Table S1). Variation partitioning among geographic distance and the dbRDA-selected environmental variables explained 30% of the variation in community composition, with environmental variables (26%) explaining much more than geographic distance (2%) (Table S2, Figure 4a). A partial-Mantel test controlling for geographic distance found that community dissimilarity significantly increased with increasing environmental dissimilarity (r = 0.19, p = < 0.001). Generalized dissimilarity modeling found that, while both spatial distance and environmental distance were significantly associated with weighted UniFrac distances, environmental distance had greater maximum predicted ecological distance and variable importance scores than geographic distance (Table S2, Figure 4b). In short, the biogeographical patterns observed for the *Bradyrhizobium* strains across the 268 soils were more closely associated with variation in environmental conditions than geographic distance per se.

**Figure 4.**
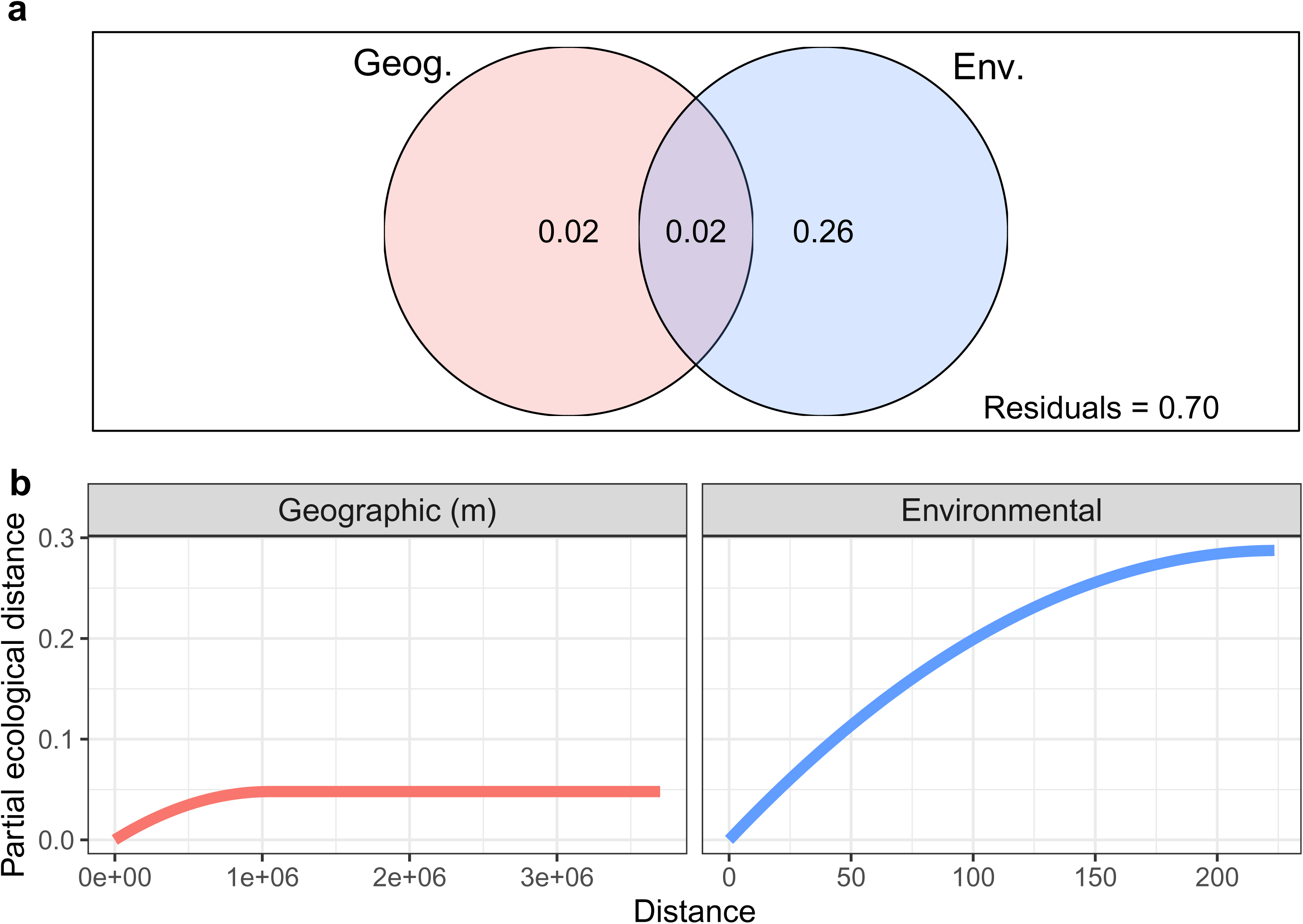
Strain-level ecological diversity of *Bradyrhizobium* across Australian soils. a) Variation partitioning of weighted UniFrac distance by geographic (latitude, longitude) and dbRDA-selected environmental variables (temperature, pH, manganese, aluminum, nitrate, and zinc). b) Partial ecological distance splines from GDM of geographic (m) and environmental (Euclidean) distance. dbRDA and GDM were performed on a subset of 219 samples to include more environmental variables.

All 181 *Bradyrhizobium* strains detected contained genes for aerobic respiration (e.g., *atpA*, *coxA*) and carbon monoxide oxidation (*coxL*) (Figure 5). Most strains also contained genes for methanol (*xoxF*), sulfide (*fccA*), and thiosulfate (*soxB*) oxidation, and the Calvin-Benson cycle (*rbcL*) for carbon fixation. There were also four stains containing genes for methane oxidation, 11 strains containing genes for the complete denitrification pathway, and 129 strains containing genes for partial denitrification. 145 strains contained *nifH* and other N-fixing genes, and 143 of those 145 also contained nodulation genes, indicating potential for symbiotic N-fixation. Only five strains contained photosynthesis genes, and those strains were not very prevalent, being detected in one to five samples (Figure 3, Figure 5).

**Figure 5.**
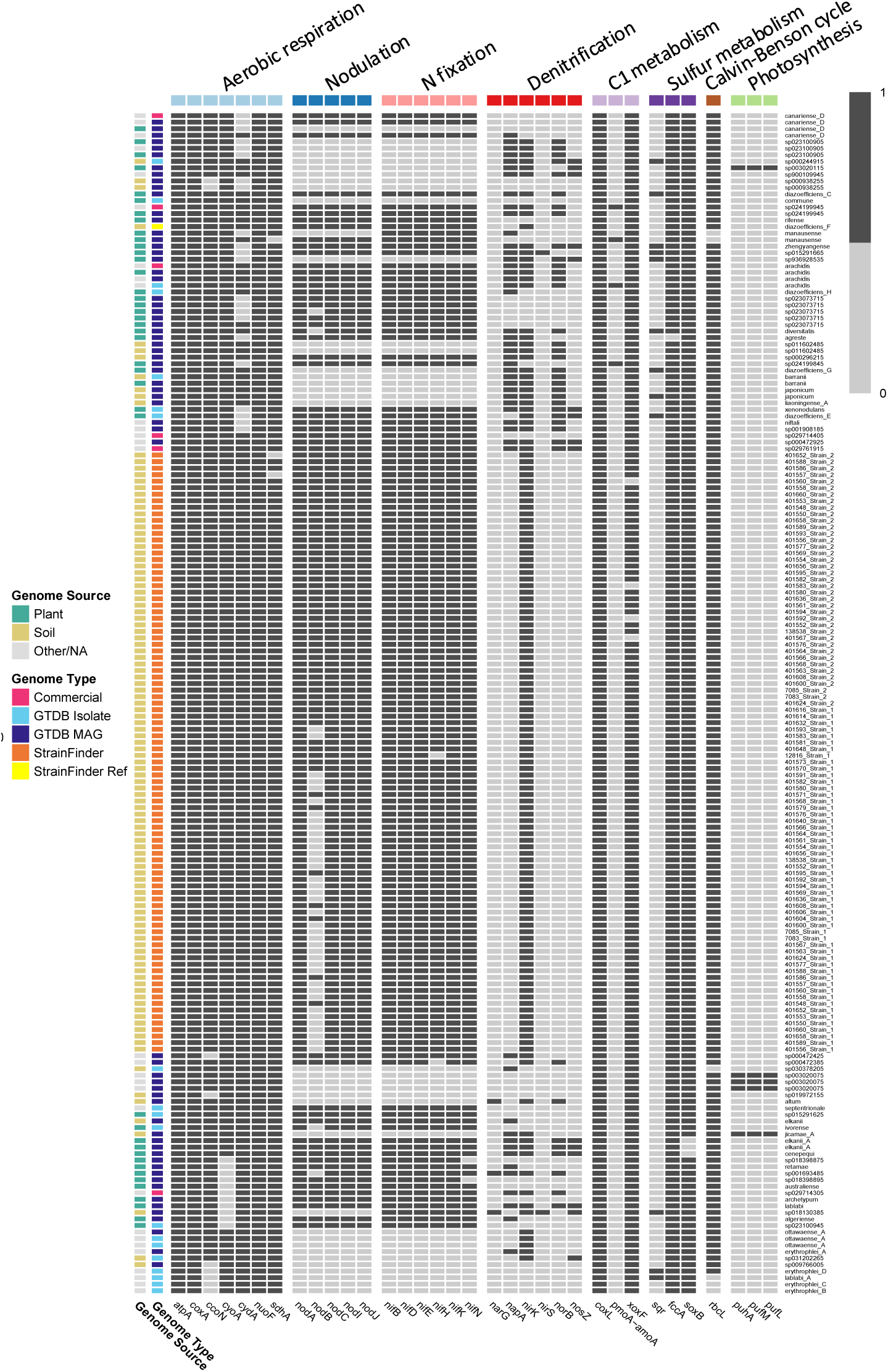
Presence or absence of genes involved in aerobic respiration, nodulation, N-fixation, C1 metabolism (carbon monoxide, methane, and methanol), sulfur metabolism (sulfide and thiosulfate), the Calvin-Benson cycle, and photosynthesis in the 181 *Bradyrhizobium* strains detected by Sylph across the 268 soil metagenomes. The ordering of the strains matches the phylogeny (Figure 3).

There were 79261 gene clusters identified in the pangenome out of 1,416,019 total genes; only a small proportion, 1935 (2%), were shared by all 181 strains, while a much larger number, 41784 (53%), were present in only one strain (Figure S10). However, the majority of gene clusters found in only one strain (73.56%) were of unknown function. Gene clusters that were present in more strains had lower geometric homogeneity than gene clusters that were present in only a few strains and this pattern was consistent across COG categories (Figure S11). Gene clusters with the lowest geometric homogeneity (i.e., greatest variability) were in the “Mobilome: prophages, transposons” COG category, followed by the “Replication, recombination and repair” categories, both of which had values < 0.85 across the gene clusters present in 10 to 172 strains (Figure S11).

## Discussion

Here we describe a generalizable methodological framework for strain-based analyses of soil bacteria that we used to conduct a comprehensive assessment of the strain-level diversity and biogeography of a ubiquitous and abundant genus of soil bacteria. We built on previous work [20, 21] by examining strain-level diversity within a genus and across a large geographic area that spanned pronounced gradients in soil and climate conditions. Our results show that there is substantial strain-level diversity in *Bradyrhizobium* across soils of the Australian continent. While we did not detect clear clustering of strains by aridity, in contrast to prior work on *Acacia*-associated *Bradyrhizobium* [8], strain-level community composition of bulk soil *Bradyrhizobium* across Australia was significantly associated with differences in environmental conditions, including mean annual temperature and soil characteristics such as soil pH and nitrate concentrations.

One reason to focus on *Bradyrhizobium* was the relatively large number of pre-existing reference genomes (compared to other soil genera). However, many of those genomes are from isolates that were cultured by growing plants in certain soils and then isolating the *Bradyrhizobium* from the nodules. Across Australian soils, we detected more MAGs (155) than isolate (26) genomes, highlighting the importance of cultivation-independent approaches for assessing the diversity of this genus in soil. We used StrainFinder to capture a broader diversity of *Bradyrhizobium* genomic diversity than is currently represented in the GTDB database. StrainFinder generated high-quality *Bradyrhizobium* MAGs based on the reference used, *B. diazoefficiens* F, which is part of the *B. japonicum* supergroup [27]. ANI values between the strains and the reference varied, meaning that StrainFinder successfully took into account the information contained in the individual soil metagenomes to generate new genomes based on the variation found in the sample, rather than just duplicating the reference genome. The StrainFinder MAGs were then detected independently by Sylph using the raw reads, including in the samples from which they were assembled, offering validation of this approach.

A total of 181 *Bradyrhizobium* strains were detected across the 331 soils, which is substantially more than the number of OTUs that would have been detected with 16S rRNA gene sequencing (Figure S8) [6]. More importantly, we were able to obtain nearly complete genomes for all of the strains, making it possible to conduct more detailed genomic analyses and make ecological inferences. Furthermore, prior work has demonstrated that 16S rRNA gene sequences alone are often unable to classify members of the genus to the species level of resolution, another benefit of our genome-resolved approach [67]. A majority of the strains detected (80%) were N-fixers, which is contrary to the finding of a dominance of free-living *Bradyrhizobium* taxa in forests from across North America [29] and a lack of *nif* and *nod* genes in many other soil *Bradyrhizobium* strains [68]. Furthermore, we only detected two *nif*-carrying free-living (i.e., lacking *nod*) *Bradyrhizobium* strains, a phenomenon which has been reported in the literature [25, 69]; in our study, 143 of 145 strains with *nif* genes also contained *nod* genes, highlighting the prevalence of symbiotic N-fixers. There were no strains that contained *nod* genes but lacked *nif* genes, suggesting a lack of “cheaters” among these 181 strains [70]. The five photosynthetic strains did not contain *nod*, consistent with previous findings [71]. In addition to N-fixation and photosynthesis, we also detected *Bradyrhizobium* strains with genes for methane oxidation and both partial and complete denitrification [72–74]. Lastly, all 181 of the Australian *Bradyrhizobium* strains contained *coxL* for carbon monoxide oxidation [54, 75] and the *xoxF* gene for methanol oxidation, consistent with previous reports [76]. Together these results highlight the diversity of energy acquisition strategies across the 181 *Bradyrhizobium* strains we detected in the Australian soils.

While agricultural areas were not included in the present study, we still detected the genomes of six out of 19 commercial *Bradyrhizobium* inoculants used in Australian agriculture [45], with one of them detected in 10 samples (Figure 3). While two (*B. canariense* D, *B. sp029714405*) of these six commercial inoculants were originally isolated from Australia, the other four were originally isolated from either South America (*B. sp024199945*, *B. sp029761915*, *B. sp029714305*) or Africa (*B. arachidis*). This raises the possibility that certain commercial agricultural inoculants have now spread into natural areas, a hypothesis in line with prior work showing that even foreign commercial inoculants have become naturalized in Australia over time [45].

The prevalence and relative abundances of *Bradyrhizobium* strains varied substantially across the Australian soils. The majority (98%) of strains were found in <10% of the soils (Figure 3), but we acknowledge that individual strains could be more prevalent across soils and our inability to detect strains may be a product of rarity (and insufficient sequencing depth), not absence per se. The strain-level biogeographic patterns observed across the genus were primarily associated with environmental dissimilarity rather than geographic distance. This result contrasts with previous evidence that *Bradyrhizobium* spp. can be dispersal limited [77]. *Bradyrhizobium* community composition was primarily associated with soil and site variables, namely temperature, pH, and concentrations of manganese, aluminum, nitrate, and zinc. Notably, aridity was not a primary variable influencing *Bradyrhizobium* community composition, contrary to our expectation based on research on forest diazotrophs [26] and *Acacia*-associated *Bradyrhizobium japonicum* strains [8]. This could be due in part to the focus of our sampling on bulk soils from a wide range of vegetation types rather than a specific focus on forests or *Acacia*-associated *Bradyrhizobium* taxa.

Mean annual temperatures across the samples studied ranged from 6.7°C to 28.1°C and in the samples with *Bradyrhizobium*, from 7.6°C to 28.1°C, highlighting their presence across the majority of the Australian temperature gradient. Some strains had a broad temperature range while others had a narrow range, and temperature was the top predictor variable of weighted UniFrac distance. However, there were no broad divisions in the phylogeny with respect to low and high temperature preferences (Figure S7). The importance of the soil and site variables is consistent with previous work on N-fixers as bacterial N-fixers as a whole functional group, including *Bradyrhizobium*, were structured more by soil properties than climate and vegetation in a Europe-wide survey [7]. In particular, that study found that relative abundances of N-fixing bacteria were greater in soils with lower pH. The importance of pH is in line with substantial previous work using amplicon sequencing of soil bacterial communities [78–80]; here we highlight, more specifically, that pH is also a key driver of strain-level variation in a ubiquitous and abundant genus.

Previous work on potential relationships between genomic traits and environmental distributions in *Bradyrhizobium* found that *Bradyrhizobium diazoefficiens* in more stressful environments (defined as environments with high temperature, low rainfall, high acidity, or high salinity) had smaller, more streamlined genomes [81]. In our study, genome size was not significantly correlated with aridity, temperature, or pH. Future work could use a landscape genomic approach similar to that employed by Simonsen (2021) [81] to assess if other genomic traits besides genome size varied across those environmental gradients. An alternative hypothesis is that genome size could also be related to prevalence or the range of environmental conditions that an organism can tolerate [82]. Our results do not support this hypothesis either, as genome size was not significantly associated with ubiquity. While it remains unclear why some strains are more ubiquitous than others and able to be detected in a larger number of soils that span a range in soil and site characteristics, the hypothesis that genome size is associated with ubiquity was not supported for members of this genus. However, it is worth noting that members of the *Bradyrhizobium* genus have larger genomes than other genera in the Bradyrhizobiaceae family, reflecting the general diversity in metabolisms and ecological strategies within the genus [28].

Given the broad spatial and environmental gradient examined here, as well as the genomic diversity encompassed by the 181 strains detected, we then asked which gene clusters in the pangenome were the most variable, in terms of presence/absence as well as geometric heterogeneity. Not surprisingly, the mobilome COG category was the most heterogeneous. However, the majority of gene clusters in each COG category were present in a low number (< 9) of strains (Figure S11). The 181 strains comprised a large pangenome of 79261 gene clusters, of which relatively few (< 3%) were in all strains, with many of the strains having unique gene clusters, similar to prior pangenomic analyses of *Bradyrhizobium* [68]. The high number of singleton gene clusters supports the idea of high genomic diversity in *Bradyrhizobium* across Australia. Previous genomic research on *Bradyrhizobium* suggested that *Bradyrhizobium* taxa contain modular systems consisting of many large integrative conjugative elements and few conjugative plasmids, which reshuffles genes and generates new combinations, which could contribute to the high number of singleton gene clusters in the pangenome [83].

## Conclusions

We sought to understand why a single bacterial genus is so ubiquitous and abundant in soil microbiomes by developing a general methodological framework for strain-level analyses of diverse microbial communities where pre-existing reference genome databases may not be sufficiently comprehensive (Figure 1). Across broad environmental gradients at a continental spatial scale, our results demonstrate that there is a strain-level phylogenetic signal in community composition associated with key soil and climatic variables. Environmental dissimilarity explained more variation in *Bradyrhizobium* community composition than spatial distance. The Australian *Bradyrhizobium* pangenome was large and dominated by singleton gene clusters, with only < 3% of gene clusters present in all 181 detected strains. The pangenome also demonstrated a breadth of metabolic strategies (Figure 5) that is also likely key to the dominance of the genus. Future work should continue to investigate strain-level environmental distributions and genomic attributes of soil bacteria, particularly in dominant taxa relevant to agriculture and key ecosystem functions.

## Acknowledgements

We acknowledge Matthew Smith for his assistance in accessing the high-quality metagenomic sequencing data and metadata from across Australia. We acknowledge the contribution of the Australian Microbiome consortium in the generation of data used in this publication.

## Author contributions

C.P.B. and N.F. conceived the overall project. C.P.B. performed analyses and drafted the manuscript. All authors provided feedback on the analyses to perform and on the interpretation of results. All authors edited the manuscript draft.()Supplementary Material sample()Supplementary material is available at the ISME Journal Online.

## Data availability

Metagenomic sequences and sample metadata are available via the Bioplatforms Australia Data Portal (https://data.bioplatforms.com/bpa/otu/metagenome). Genome sequences for the 181 detected *Bradyrhizobium* genomes are available on figshare (10.6084/m9.figshare.29176142). Downstream data, including the list of sample IDs, and analysis scripts are available on Zenodo (10.5281/zenodo.15540063).

