## Supplemental Figures S1-S11 for "Continent-wide assessment of the strain-level diversity of *Bradyrhizobium*, a dominant soil bacterial genus"

Running head: *Bradyrhizobium* strain diversity and ecology

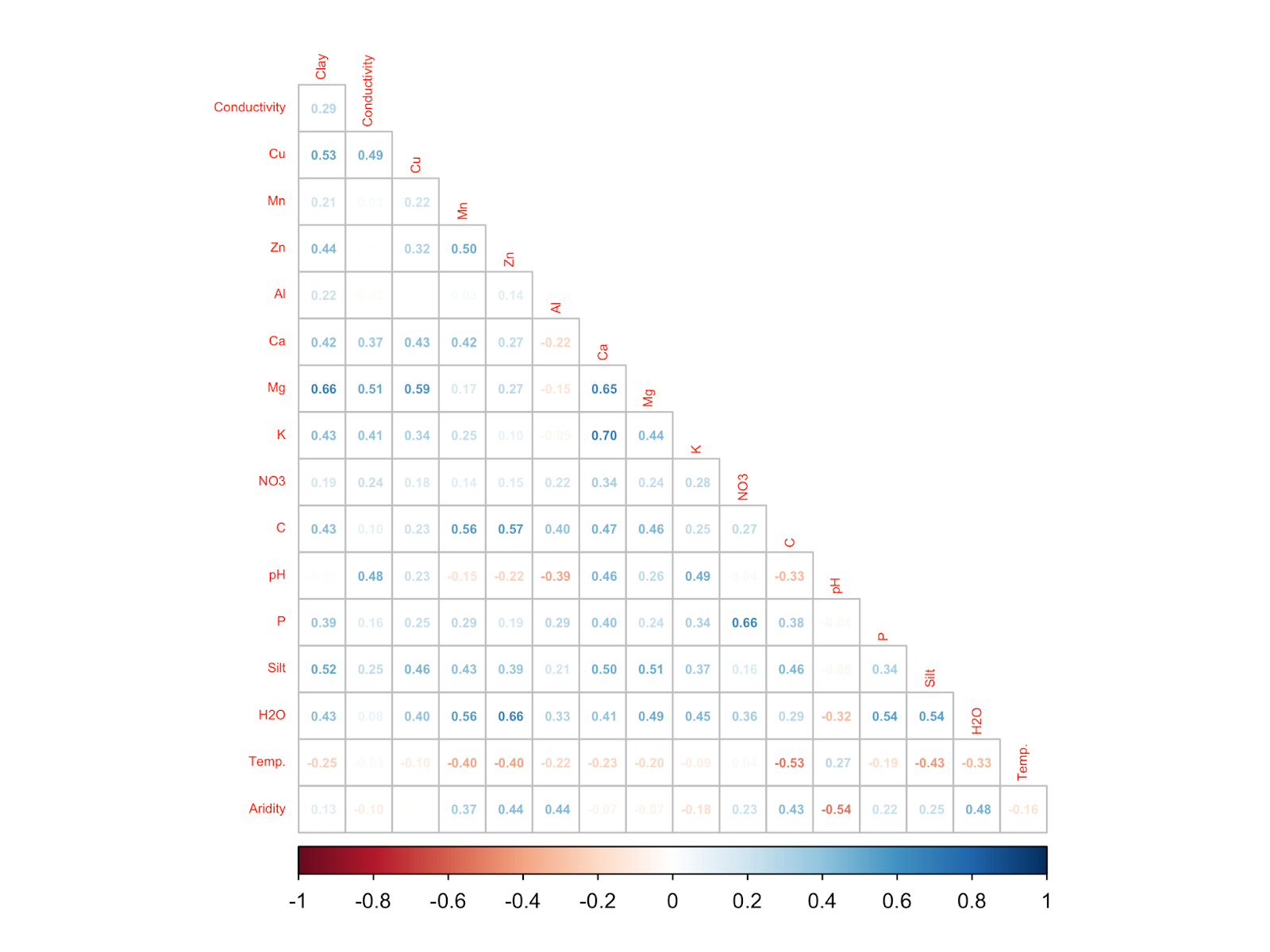

Figure S1. Pearson correlations among environmental variables across the 331 samples. Boron, sodium, mean annual precipitation, % sand, and iron were excluded from the community-level analyses as they had correlations > 0.7 with at least one other variable.

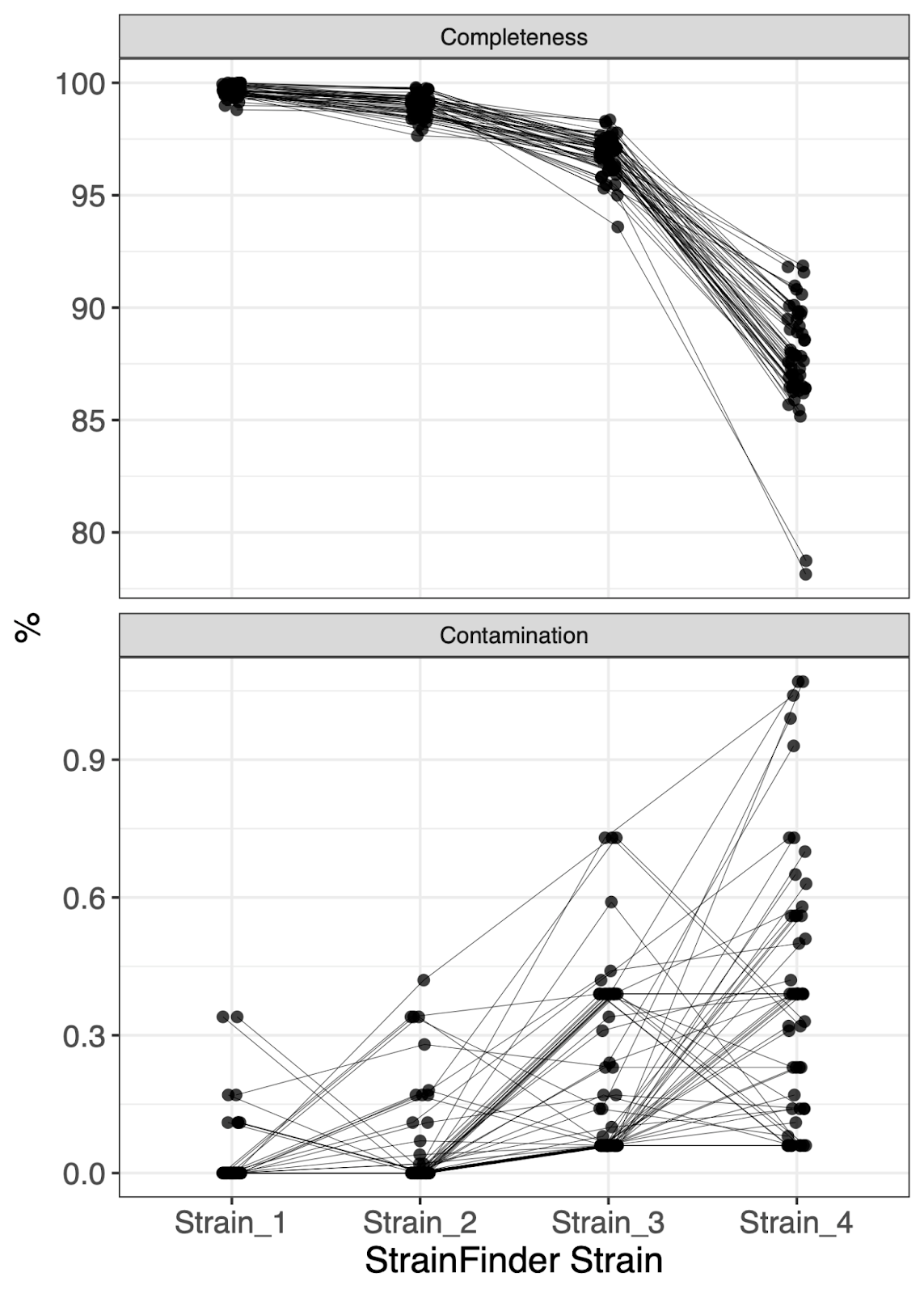

Figure S2. Completeness and contamination of *Bradyrhizobium* MAGs assembled using StrainFinder. Only the first two (the most abundant) *Bradyrhizobium* genomes per metagenome were used in downstream analyses. Lines connect strains from the same sample (n = 4 per sample).

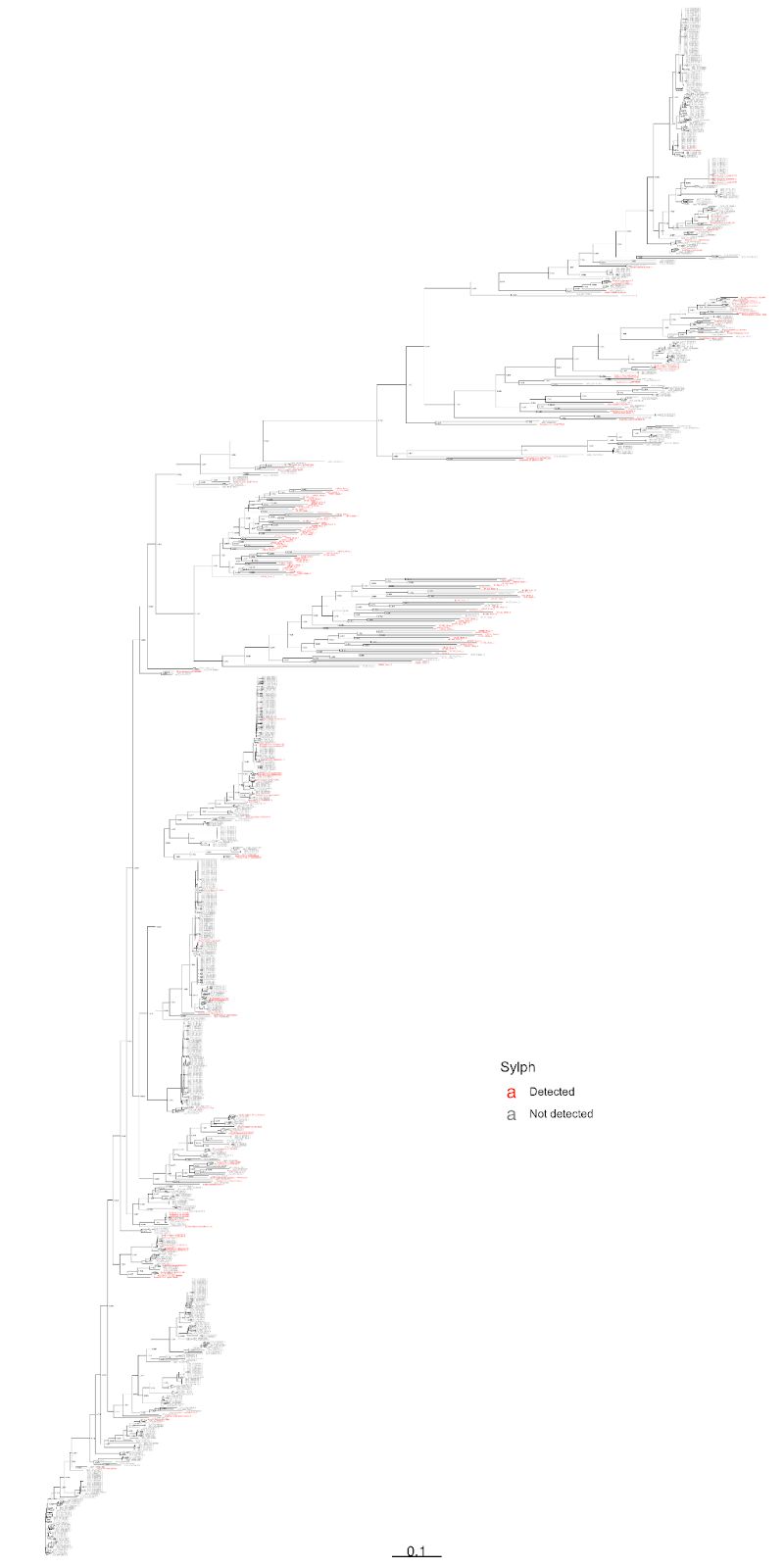

Figure S3. Phylogenetic tree of 915 *Bradyrhizobium* genomes (106 StrainFinder MAGs, 809 GTDB genomes) based on a concatenated alignment of 2409 amino acids from the bac120 marker gene set. The tree shows that the 181 strains detected by Sylph in the Australian soil metagenomes are distributed throughout the tree.

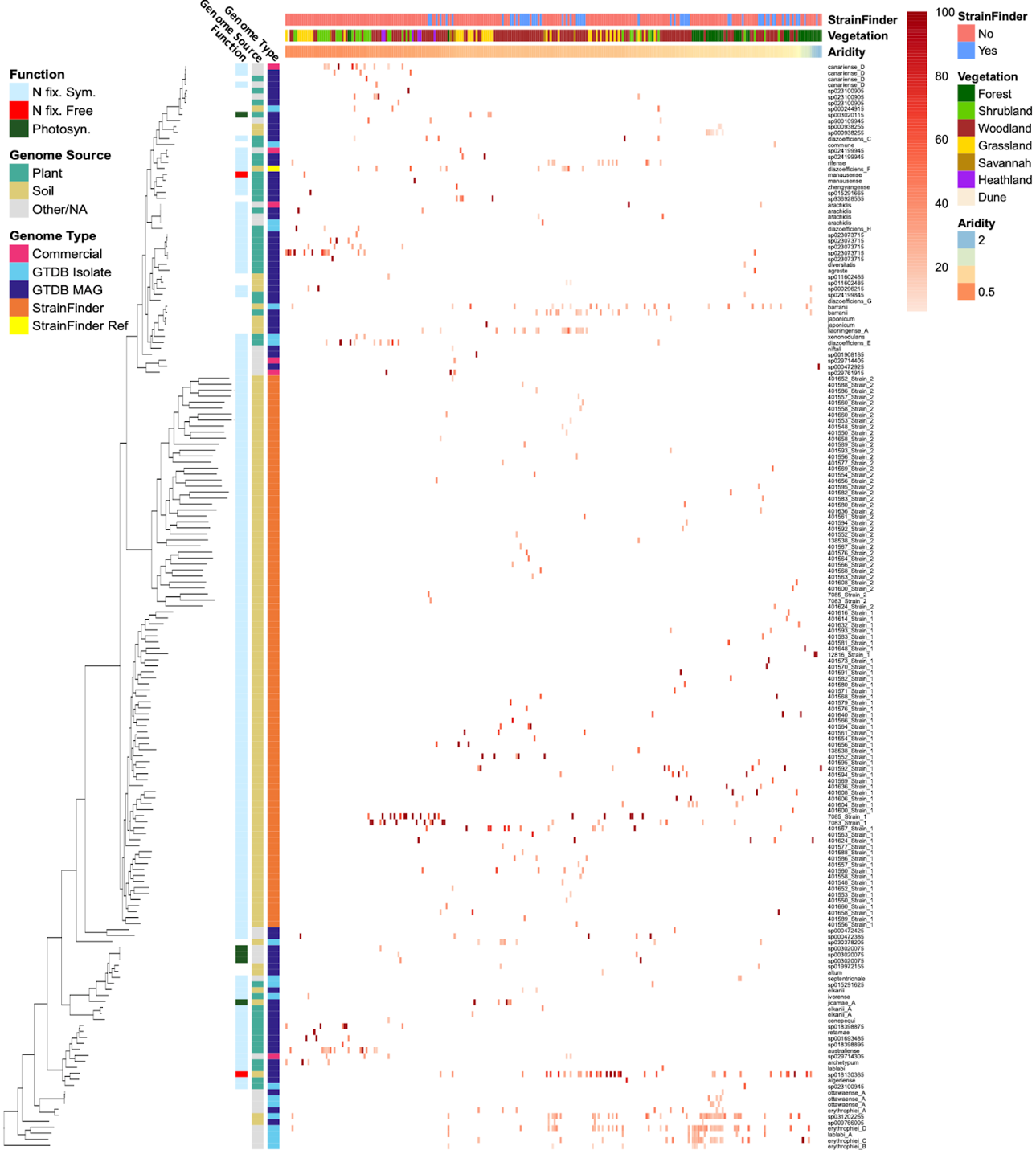

Figure S4. *Bradyrhizobium* percent relative abundances from Sylph run on 331 samples and 915 *Bradyrhizobium* genomes (809 GTDB genomes + 106 StrainFinder MAGs). We note that this figure is identical to Figure 3, except relative abundances of each strain per soil metagenome are indicated. Shown are 181 strains (89 GTDB + 92 StrainFinder) detected in at least one of the 331 samples (shown are 268 samples with at least one *Bradyrhizobium* strain detected). Columns (samples) are sorted by increasing aridity index from left to right while rows (genomes) are sorted according to the phylogenetic tree shown on the left, based on a concatenated alignment of 2026 amino acids from the bac120 marker gene set. Shown above the heatmap (legend on right) are sample metadata (whether or not StrainFinder was run on the sample, vegetation type, and aridity index). Shown to the left of the heatmap (legend on left) are genome metadata (function, genome source, and genome type). Genome source “Other/NA” means that the source was not stated in the metadata.

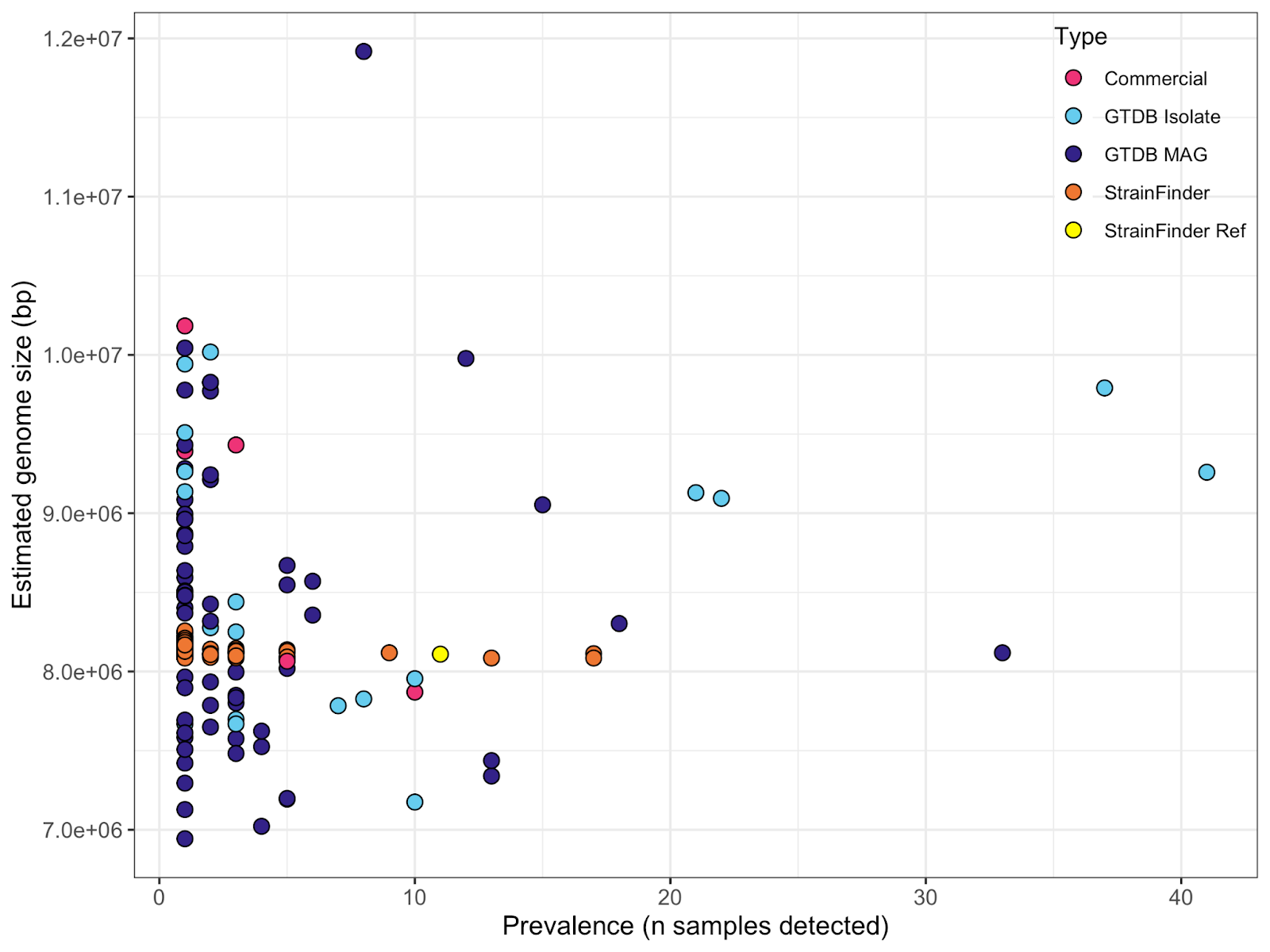

Figure S5. Lack of significant relationship between prevalence (number of soil metagenomes in which a given strain was detected) and estimated genome size for the 181 *Bradyrhizobium* strains detected by Sylph (R^2^ = 0.02, p = 0.06).

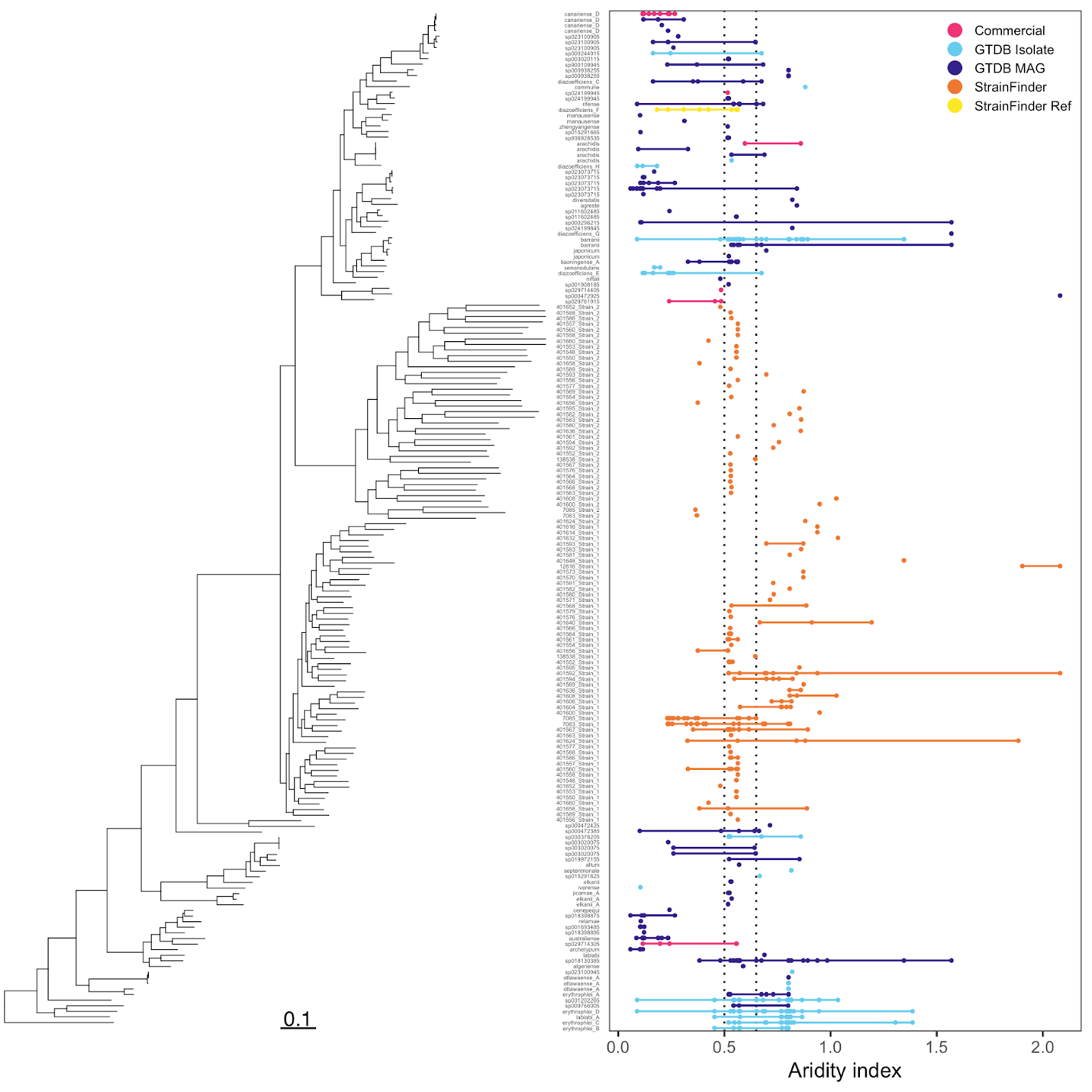

Figure S6. Phylogenetic tree of 181 *Bradyrhizobium* strains detected by Sylph across 268 soil metagenomes based on a concatenated alignment of 2026 amino acids from the bac120 marker gene set, as well as the aridity index values of the samples in which they were detected. The tree is rooted with *Nitrobacter winogradsky* (not shown). Dotted lines show the cutoffs for climate classes where arid to semi-arid is < 0.5, dry sub-humid is 0.5 to 0.65, and humid is > 0.65.

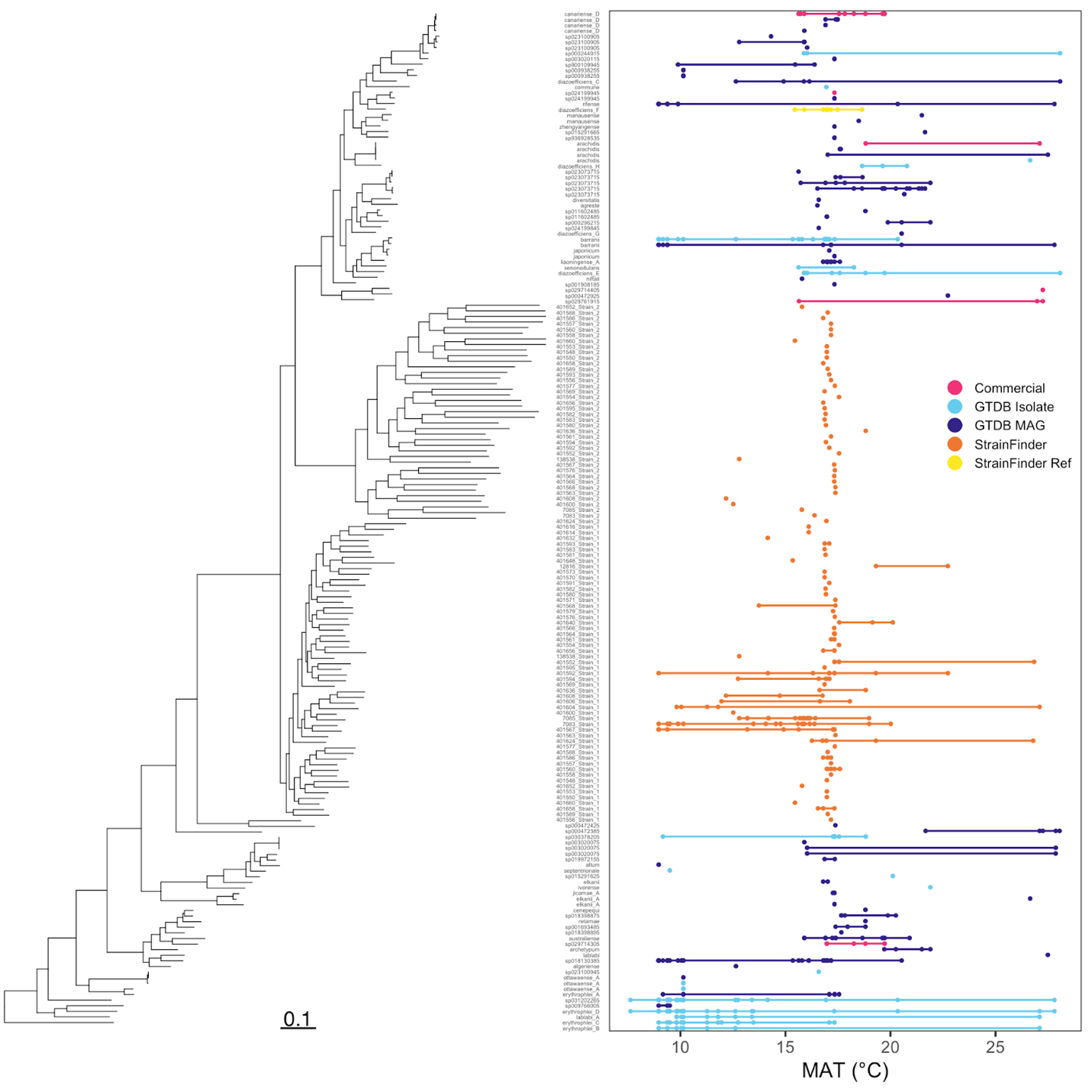

Figure S7. Phylogenetic tree of 181 *Bradyrhizobium* strains detected by Sylph across 268 soil metagenomes based on a concatenated alignment of 2026 amino acids from the bac120 marker gene set, as well as the mean annual temperature of the samples in which they were detected. The tree is rooted with *Nitrobacter winogradsky* (not shown).

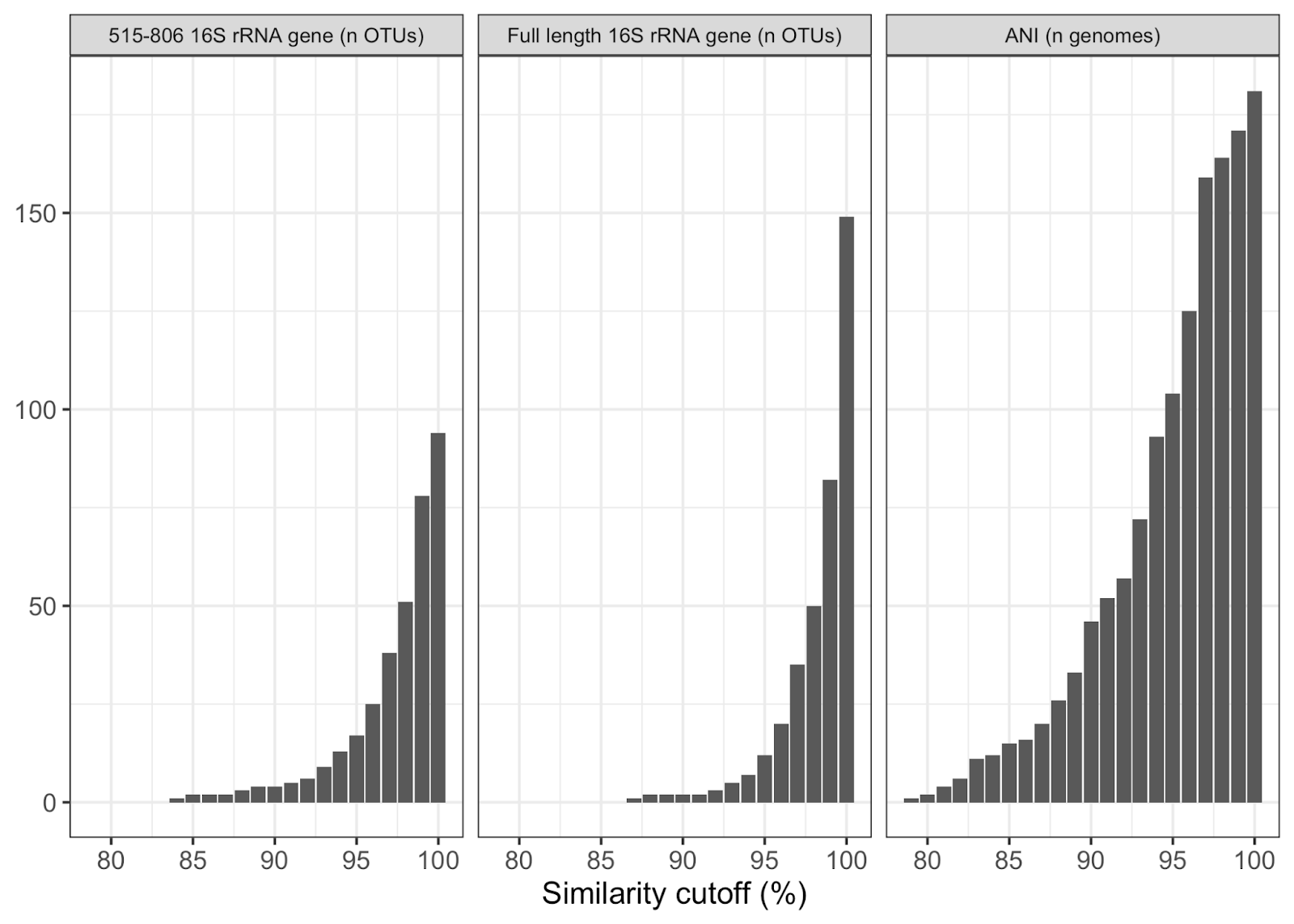

Figure S8. The number of operational taxonomic units (OTUs, left and center) and number of genomes (right) that the 181 strains would be clustered into using V4 (positions 515-806) 16S rRNA gene sequences (left), full length 16S rRNA gene sequences (center), or whole genome average nucleotide identity (ANI, right) similarity cutoffs. Note that two strains did not contain the full length 16S rRNA gene and were omitted from the 16S rRNA gene analysis shown here.

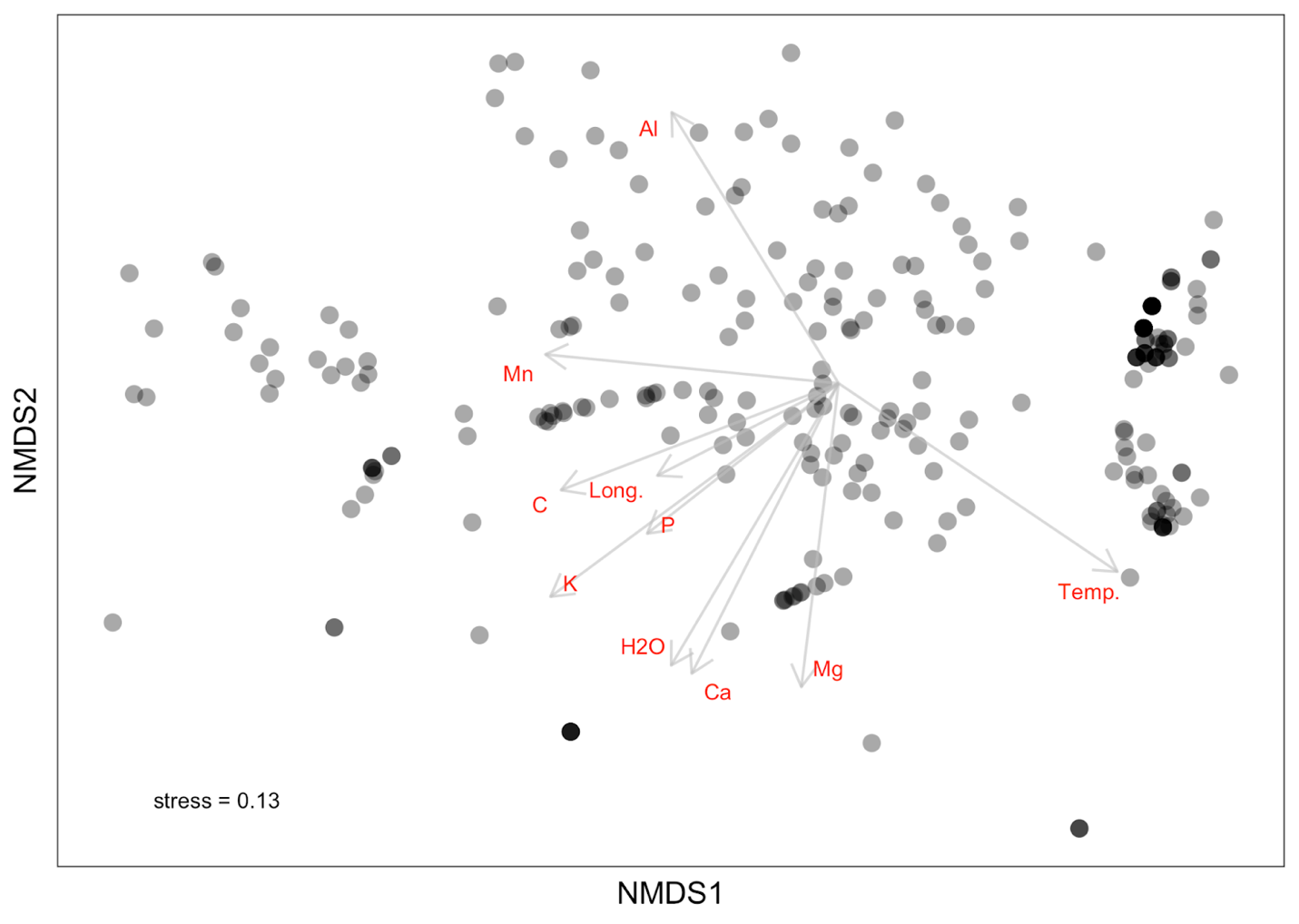

Figure S9. Non-metric multidimensional scaling (NMDS) of weighted UniFrac distance (stress = 0.13), with highly and significantly associated environmental variables (envfit, p = 0.001) overlaid as vectors. Latitude, copper, zinc, percent silt, and pH (not shown) were also significantly associated with the ordination (envfit, p < 0.05). Temp. = mean annual temperature, Al = aluminum, Mn = manganese, P = Colwell phosphorus, Long. = longitude, H2O = water content, C = % organic carbon, K = potassium, Ca = calcium, Mg = magnesium. The sample size shown here is 331, and some environmental variables contain NAs (maximum number of NA values is water content, with 67 NA).

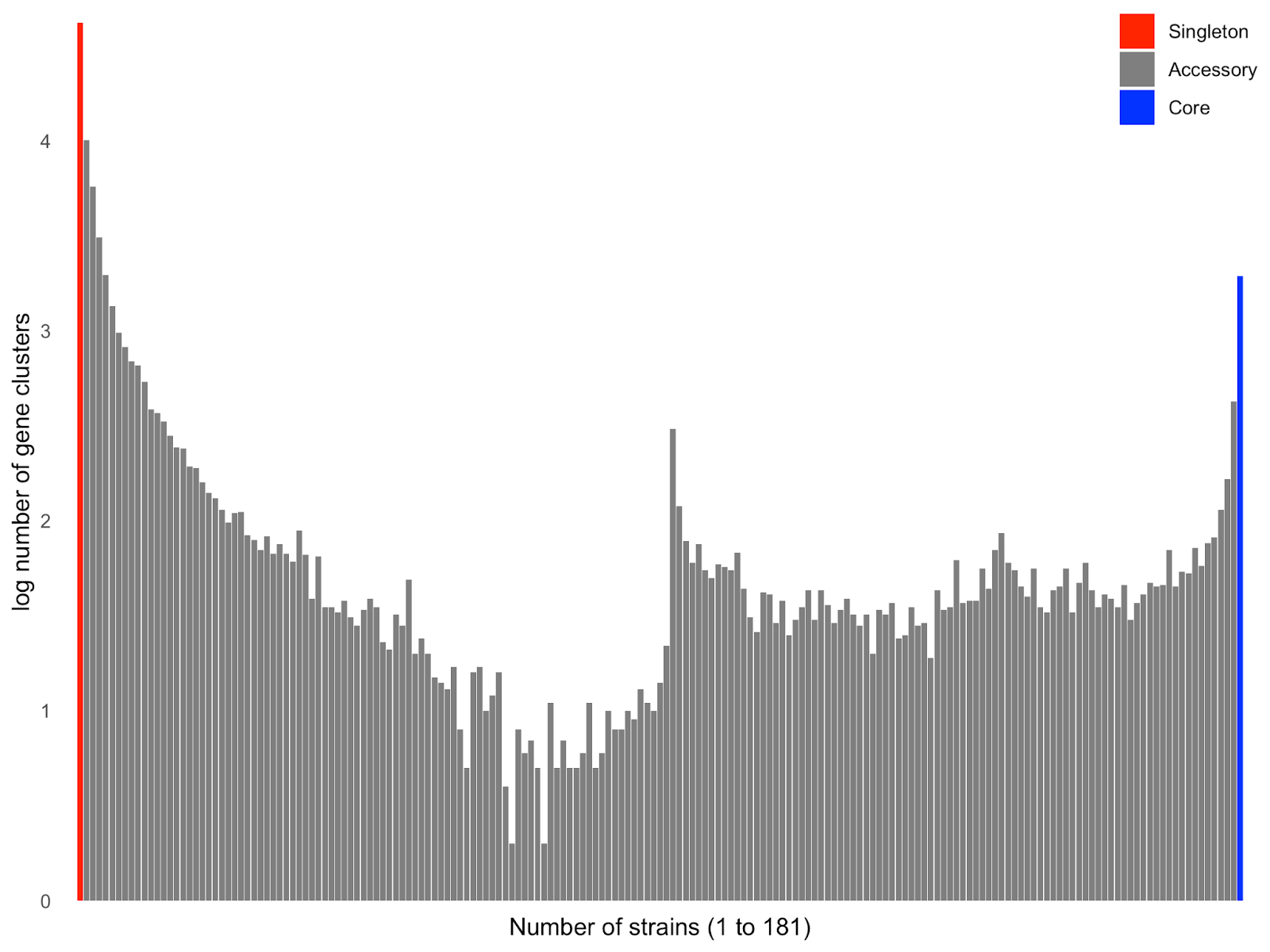

Figure S10. Number of gene clusters (log transformed) as a function of how many strains they were found in. Number of singleton gene clusters = 41784, number of accessory gene clusters (i.e., clusters found in 2 to 180 strains) = 35542, number of core gene clusters (i.e., clusters shared across all 181 strains) = 1935.

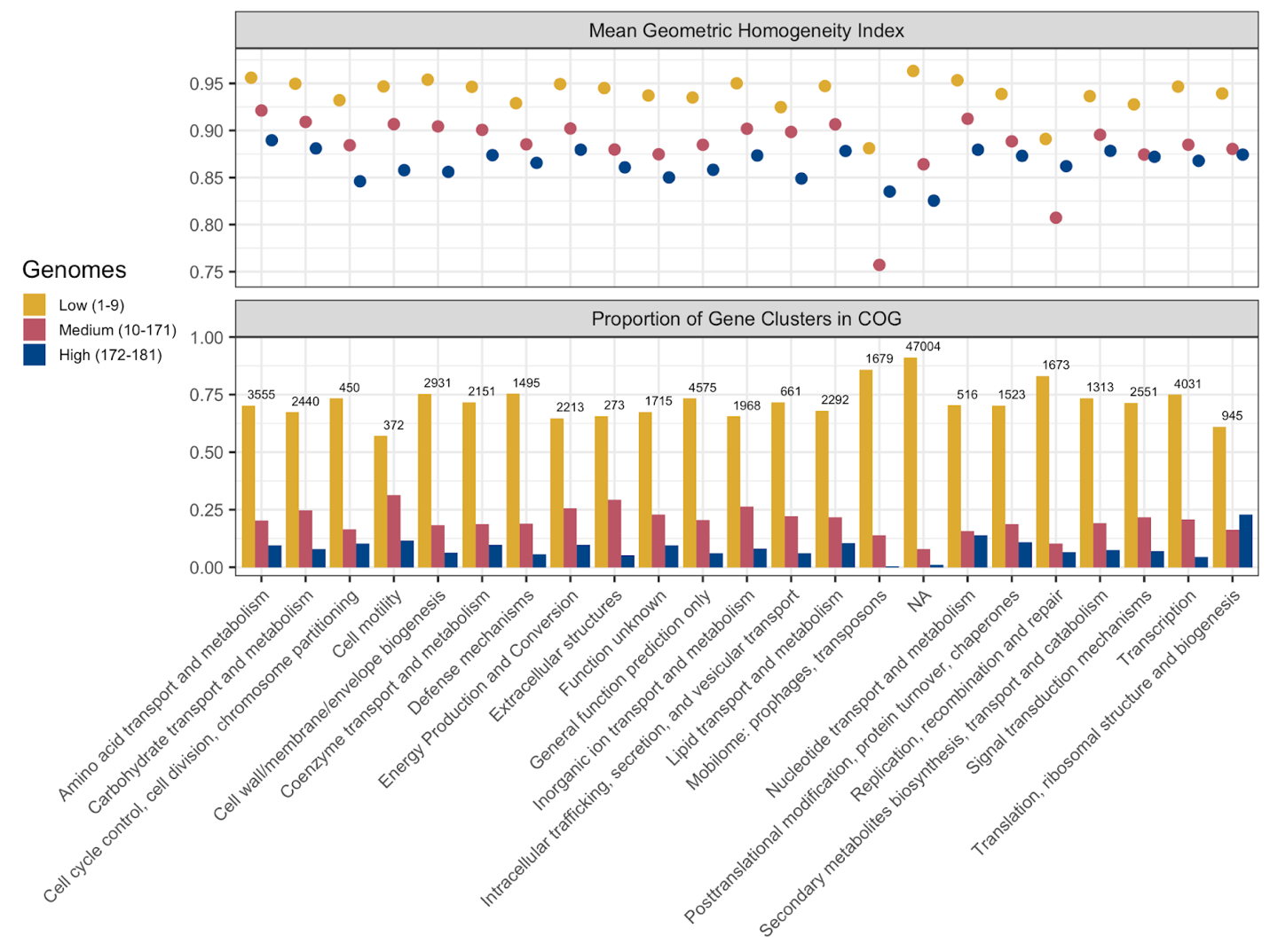

Figure S11. Mean geometric homogeneity values and proportion of gene clusters in each COG category, grouped by how many genomes the gene clusters were found in (low = 1-9 genomes, medium = 10-171 genomes, high = 172-181 genomes). The 9 and 172 cutoffs correspond to 5% and 95% of the genomes. Most gene clusters in each COG were found in 9 or fewer genomes. Numbers above the bars represent the total number of gene clusters in each COG. NA = no COG category assigned (i.e., unannotated).
